# Long-term dysbiosis promotes insulin resistance during obesity despite rapid diet-induced changes in the gut microbiome of mice

**DOI:** 10.1101/116095

**Authors:** Kevin P. Foley, Emmanuel Denou, Brittany M. Duggan, Rebecca Chan, Jennifer C. Stearns, Jonathan D. Schertzer

**Author notes:** **Corresponding author**: Dr. Jonathan D. Schertzer, Department of Biochemistry and Biomedical Sciences, Faculty of Health Sciences, McMaster University, HSC 4H30D; 1200 Main Street West, Hamilton, Ontario, Canada, L8N 3Z5.

## Abstract

The intestinal microbiota and insulin sensitivity are rapidly altered in response to a high fat diet (HFD). It is unclear if gut dysbiosis precedes insulin resistance or vice versa. The initial triggers of diet-induced insulin resistance can differ from mechanisms underlying chronic dysglycemia during prolonged obesity. It is not clear if intestinal dysbiosis contributes to insulin resistance during short-term or long-term HFD-feeding. We found that diet-induced changes in the composition of the fecal microbiome preceded changes in glucose and insulin tolerance at both the onset and removal of a HFD in mice. Dysbiosis occurred after 1-3 days of HFD-feeding, whereas insulin and glucose intolerance manifested by 3-4 days. Antibiotic treatment did not alter glucose tolerance during this short-term HFD period. Conversely, antibiotics improved glucose tolerance in mice with protracted obesity caused by long-term HFD feeding for over 2 months. We also found that microbiota transmissible glucose intolerance only occurred after prolonged diet-induced dysbiosis. Germ-free mice had impaired glucose tolerance when reconstituted with the microbiota from long-term, but not short-term HFD-fed animals. Our results are consistent with intestinal microbiota contributing to chronic insulin resistance and dysglycemia during prolonged obesity, despite rapid diet-induced changes in the taxonomic composition of the fecal microbiota.

---

Type 2 diabetes (T2D) is characterized by dysglycemia, which can manifest from insulin resistance and insufficient insulin secretion. Insulin resistance can last for many years and the majority of individuals with features of prediabetes, such as glucose intolerance, eventually develop T2D^1^. Environmental factors such as diet and exercise contribute to the increased prevalence of prediabetes, which coincides with increased incidences of obesity^2^. The composition of the intestinal microbiota can also influence postprandial glucose responses^3^. This adds microbes to the list of environmental factors that can influence glucose tolerance and possibly T2D risk. The composition of the gut microbiota is altered in obesity, T2D and by insulin sensitizing drugs^4–6^. However, it is not clear how the microbiota contributes to the progression of glucose intolerance versus obesity. This is an important distinction because not all obese individuals develop insulin resistance, glucose intolerance or T2D. Further, the timing and progression of obesity can be different from and dysglycemia.

The intestinal microbiota can contribute to host energy balance and lipid deposition through hormonal cues^7^. Gut dysbiosis during obesity is sufficient to increase adiposity, independently of host genetics^8,9^. Hence, there is a strong connection between the microbiota and obesity, but it is not clear if a microbiota-induced change in adiposity is the primary factor contributing to glucose intolerance. Increased adiposity correlates with glucose intolerance, but there may be other microbiota-driven mechanisms that alter blood glucose and insulin sensitivity. Microbes can influence circulating metabolites such as branched chain amino acids and consequently alter insulin sensitivity^10^. In addition, microbial components can subvert the intestinal or other mucosal barriers to promote inflammation and glucose intolerance^11,12^. Conversely, microbiota transfer from lean donors can increase peripheral insulin sensitivity in obese patients, independent of changes in adiposity^13^.

Diet is a key factor in the development of obesity and a dominant factor shaping the composition of the microbiota. In mice, diet influences the composition of the gut microbiome more than host genetics and it is already known that increasing dietary fat content can rapidly perturb the composition of the microbiota within days^14^. A high fat diet (HFD) induces obesity, insulin resistance and glucose intolerance in rodents. However, the mechanisms underpinning glucose intolerance during the initial stages of HFD feeding are different than those governing chronic glucose intolerance during prolonged HFD-induced obesity. For example, ectopic lipid accumulation in the skeletal muscle and liver is associated with glucose intolerance after the first few days of HFD feeding, whereas metabolic tissue inflammation plays a more prominent role in propagating glucose intolerance after months of HFD-feeding in mice^15^. It is not clear if HFD-induced changes in the microbiota contribute to the mechanisms underpinning acute versus chronic insulin resistance and glucose intolerance.

We found that HFD-induced changes in the composition of the microbiota preceded overt dysglycemia in mice. Hence, we hypothesized that dysbiosis caused by both short-term and long-term HFD-feeding would promote increased fat accumulation and glucose intolerance. However, we found that only long-term HFD-feeding promoted transmissible glucose intolerance, which can occur independently of changes in adiposity.

## Results

### Short-term HFD feeding causes glucose intolerance nce in mice

Glucose tolerance was not different after 1 or 2 days of HFD-feeding (Fig. 1A, S1A). Insulin intolerance was evident after 3 days of feeding mice a 60% HFD, but not 3 days of 45% HFD (Fig. S1B). Four days of HFD-feeding marked the first time that both 45% and 60% HFD caused glucose intolerance compared to mice fed a chow diet (Fig. 1B). Glucose intolerance persisted after 14 days (Fig. 1C) and 14 weeks of feeding 45% or 60% HFD (Fig. 1D). Insulin intolerance was evident after 7 days and 12 weeks of either 45% or 60% HFD-feeding (Fig. S1C, D). The 60% HFD caused higher glucose intolerance and higher insulin intolerance compared to the 45% HFD, when tested between 4 days and 14 weeks of HFD-feeding (Fig. 1, S1). These results show that it takes 3-4 days of HFD feeding to cause glucose and insulin intolerance in mice. These results also show that the percentage of dietary fat is directly related to the level of glucose and insulin intolerance in mice.

**Figure 1:**
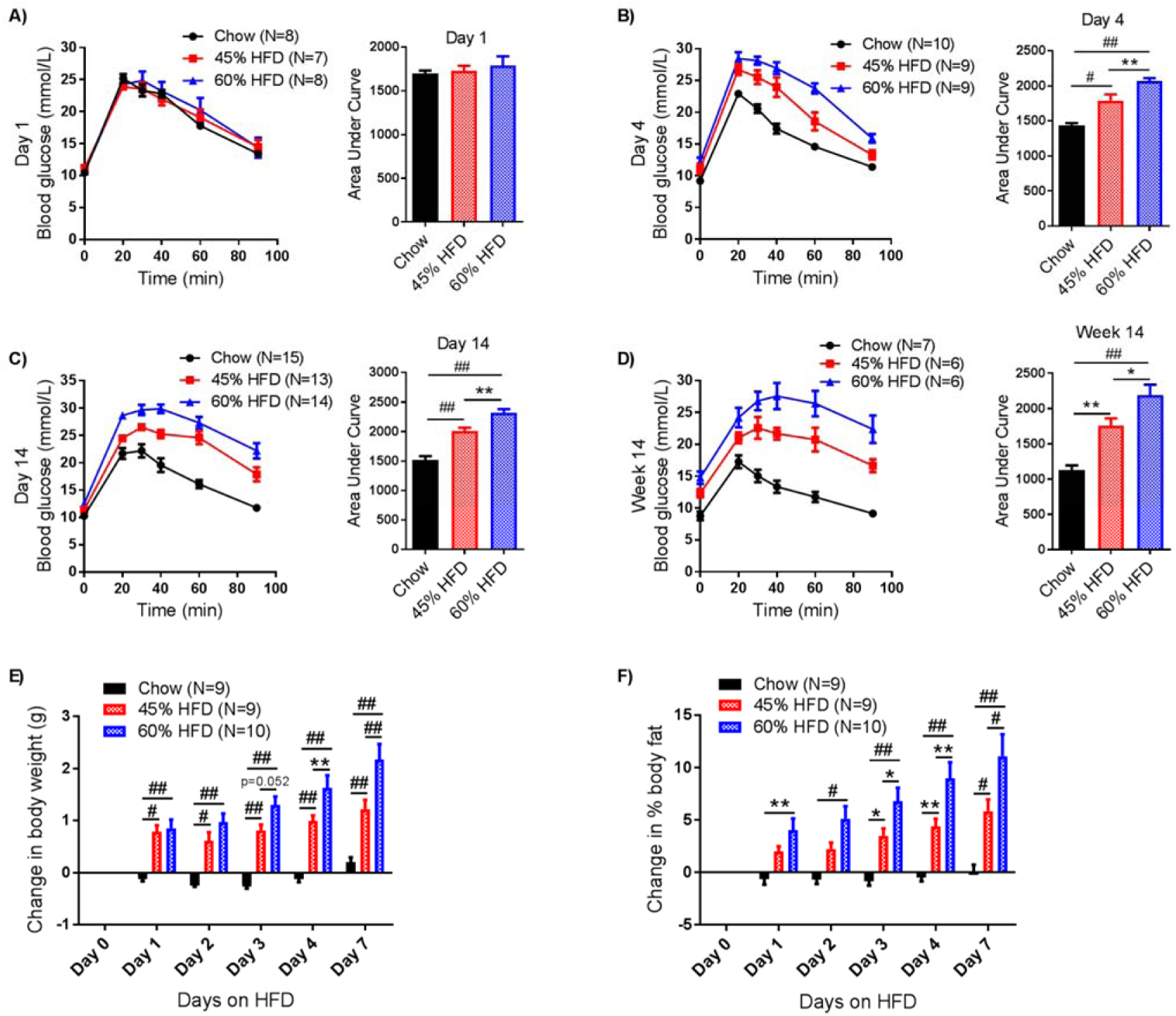
High fat feeding for 4 days is sufficient to induce glucose intolerance and increase adiposity in mice. A-D) Mice were fed a chow, 45% HFD, or 60% HFD for 1 day (A: N=8, 7, 8), 4 days (B: N=10, 9, 9), 14 days (C: N=15, 13, 14), or 14 weeks (D: N=7, 6, 6) before being tested for glucose tolerance with a 2 g/kg (A-C) or 0.9 g/Kg (D) dose by *i.p.* injection. Blood glucose measures were taken at indicated time points. Each figure shows the GTT/ITT curve and AUC. Statistical significance was measured as p<0.05 using one-way ANOVA. Post Hoc analysis was performed using Tukey’s multiple comparisons test (* p<0.05; ** p<0.01; # p<0.001; ## p<.0001). E-F) Body weight and adiposity were measured in mice fed a chow, 45% HFD, or 60% HFD for 7 days (N= 9, 9, 10). Adiposity was measured as % body fat using whole body echo-MRI imaging. Change in body weight (E) and change in % body fat (F) were calculated as the difference between Day 0 and Day “X” measures within each animal. Statistical significance was measured as p<0.05 using two-way ANOVA with repeated measures (time). Post Hoc analysis was performed using Tukey’s multiple comparisons test (* p<0.05; ** p<0.01; # p<0.001; ## p<.0001)

### Increased adiposity precedes glucose intolerance HFD-fed mice

A single day of feeding either 45% or 60% HFD increased body mass compared to chow-fed mice (Fig. 1E). Between 4-7 days of HFD-feeding, a 60% HFD caused a greater change in body mass compared to a 45% HFD (Fig. 1E). A single day of feeding 60% HFD increased adiposity (i.e. body fat percentage), whereas it took 3 days of feeding a 45% HFD to detect an increase in whole body adiposity (Fig. 1F). Between 3-7 days of HFD feeding, a 60% HFD caused a greater increase in adiposity compared to feeding mice a 45% HFD (Fig. 1F). These results show that: 1) a single day of HFD-feeding increased body mass and body fat percentage and 2) the dietary fat content is directly related to the level of adiposity during the first week of HFD-feeding.

### HFD-induced changes in the microbiome precede glucose intolerance in mice

We next assessed the timing of changes in the taxonomy and predicted metagenomic characteristics of the fecal microbiome during the first week of HFD-feeding. The Bray-Curtis similarity index of fecal bacterial DNA showed that the bacterial community was different in chow fed mice (i.e. day 0), compared to mice fed a 45% or 60% HFD for 7 days (Fig. 2A, B). The similarity matrix showed a progressive increase between day 1 and day 4 of feeding a 45% HFD diet (Fig. 2A). Only 1 day of 60% HFD-feeding equated to Bray-Curtis similarity index that was similar to 7 days on the 60% HFD diet (Fig. 2B). Supplemental Figure 2 shows phylum level changes in the feces of mice when fed chow, 45% or 60% HFD for up to 98 days (i.e. 14 weeks). The Firmicutes/Bacteroidetes ratio was increased by 2-3 days of HFD-feeding, where the 60% HFD caused a larger magnitude of change in this ratio (Fig. 2C).

**Figure 2:**
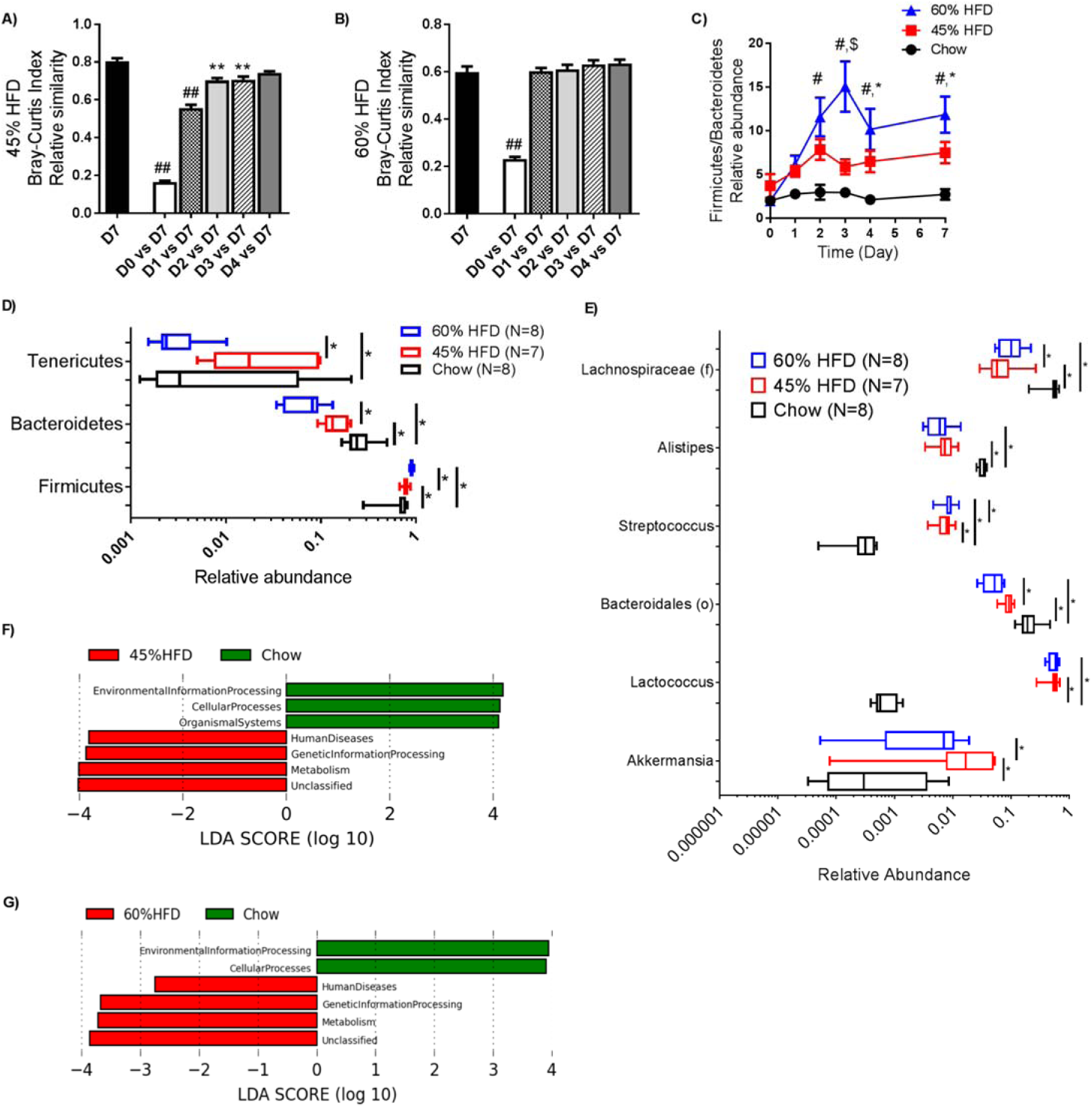
High fat feeding induces changes in the fecal microbiota that precede glucose intolerance in mice. Fecal samples were taken over the first 7 days of high fat feeding and processed for bacterial DNA sequencing (N=7-14). Beta-diversity was measured based on the Bray-Curtis dissimilarity index relative to the microbial profile at day 7 in mice fed with 45% (A) or 60% HFD (B) from Day 0 to 7. C) The Firmicutes/Bacteroidetes ratio over the first 7 days of high fat feeding. Phylum (D) and Genus (E) level changes in the microbiome on Day 3 of high fat feeding with 45% or 60% HFD. LDA scores from LEfSe analysis of PICRUSt-predicted function 3 days after 45% (F) and 60% (G) HFD relative to chow diet. ANOVA was used for all figures and false discovery rate was accounted for by controlling the familywise error rate using the Bonferroni correction. Significance was accepted at p<0.05. Panel A and B: * p<0.05; ** p<0.01; # p<0.001; ## p<.0001). Panel C: “*” shows differences between chow and 45%; “#” between chow and 60% and “$” between 45% and 60% HFD profiles.

We then focussed on the 3^rd^ day of HFD-feeding, since this time-point preceded overt glucose and insulin intolerance due to both 45% and 60% HFD. Three days of feeding mice either a 45% or 60% HFD equated to higher relative abundance of Firmicutes and lower abundance of Bacteroidetes (Fig. 2D). At the genus level, 3 days of feeding either HFD caused a striking increase in the relative abundance of *Lactococcus* and *Streptococcus* (Fig. 2E). The relative abundance of Lachnospiraceae (f), *Alistipes,* and Bacteroidales (o) were decreased after 3 days of feeding of either HFD (Fig. 2E). *Akkermansia* was higher after 3 days of feeding a 45% HFD (Fig. 2E). Many aspects of taxonomic changes observed on Day 3 of HFD feeding were maintained at later time points of HFD feeding in mice (Fig. S2, S3).

Using phylogenetic investigation of communities by reconstruction of unobserved states (PICRUSt), our results showed that short-term HFD-feeding altered the predicted metagenome and that a 45% and 60% HFD had similar effects. In comparison to the feces from chow-fed mice, LDA (Linear Discriminant Analysis) Effect Size (LEFSe) analysis resulted in significant LDA scores for environmental processing, cellular processes, human diseases and metabolism after 3 days of feeding 45% HFD or 60% HFD (Fig. 2F, G). We previously showed that 12 weeks of HFD-feeding decreased the PICRUSt-predicted genetic capacity of the fecal microbiota for metabolism and increased the predicted genetic capacity related to environmental information processing^16^. Here, our results after LEfSe analysis show that 3 days of feeding either a 45% or 60% HFD promoted discordant effects on PICRUSt-predicted metabolism and environmental information processing pathways in the fecal microbiota (Fig 2F, G). Specifically, we found that 3 days of HFD feeding caused an increase in the predicted genetic capacity for metabolism and a decrease in environmental information processing (Supplemental Table 1). Therefore, short-term HFD-induced effects on predicted genetic capacity for metabolism and environmental information processing were opposite to those during long-term HFD feeding.

We next analyzed glucose tolerance and microbiota composition after short-term removal of the HFD and replacement with a chow diet by feeding mice a 60% HFD for 14 days, then switching the mice to a chow diet for 2 days (i.e. Day 16) (Figure 3A). Despite HFD removal, the mice previously fed 60% HFD remained glucose intolerant and still had higher body mass when compared to age-matched chow-fed mice (Fig. 3B, C). The Bray-Curtis similarity matrix showed that the bacterial community was different 2 days after removing a 60% HFD (Fig. 3D). When compared to 14 days of HFD-feeding, this similarity matrix showed that only 2 days of returning mice to a chow diet after 14 days of HFD-feeding resembled the difference seen in mice continually fed a chow diet for 14 or 16 days (Fig. 3D). Average phylum level changes in the feces of mice fed a chow diet or a 60% HFD and removal back to chow for 2 days are shown in Figure S3A. At the phylum level, 14 days of HFD-feeding equated to a higher Firmicutes/Bacteroidetes ratio compared to chow-fed mice, but replacing the HFD with a chow diet for 2 days resulted in a lower Firmicutes/Bacteroidetes ratio to a level that was similar to mice continually fed a chow diet for 14 or 16 days (Fig. 3E). Removing the HFD for 2 days also restored HFD-induced changes in the relative abundance of Firmicutes and Bacteroidetes, but not Verrucomicrobia (Fig. 3F). At the genus level, many of the same changes observed after 7 days of HFD were also evident after 14 days of a 60% HFD (Fig. 3G, S3B). A notable exception was *Akkermanisia,* which was lower after 14 days of 60% HFD-feeding (Fig. 3G). Replacing the HFD with a chow diet for 2 days reduced the abundance of *Lactococcus* and increased members of the order Bacteroidales to levels observed in chow-fed controls (Fig. 3G). Genus-level changes over the first 2 weeks of feeding a 45% or 60% HFD, plus removal of the HFD for 2 days are shown in Supplemental Figure 3B. Overall, these data show that diet-induced changes in fecal microbiome precede changes in glucose and insulin intolerance at both the onset and removal of a HFD. These data also show that the magnitudes of change in the microbiome are directly related to the fat content of the diet.

**Figure 3:**
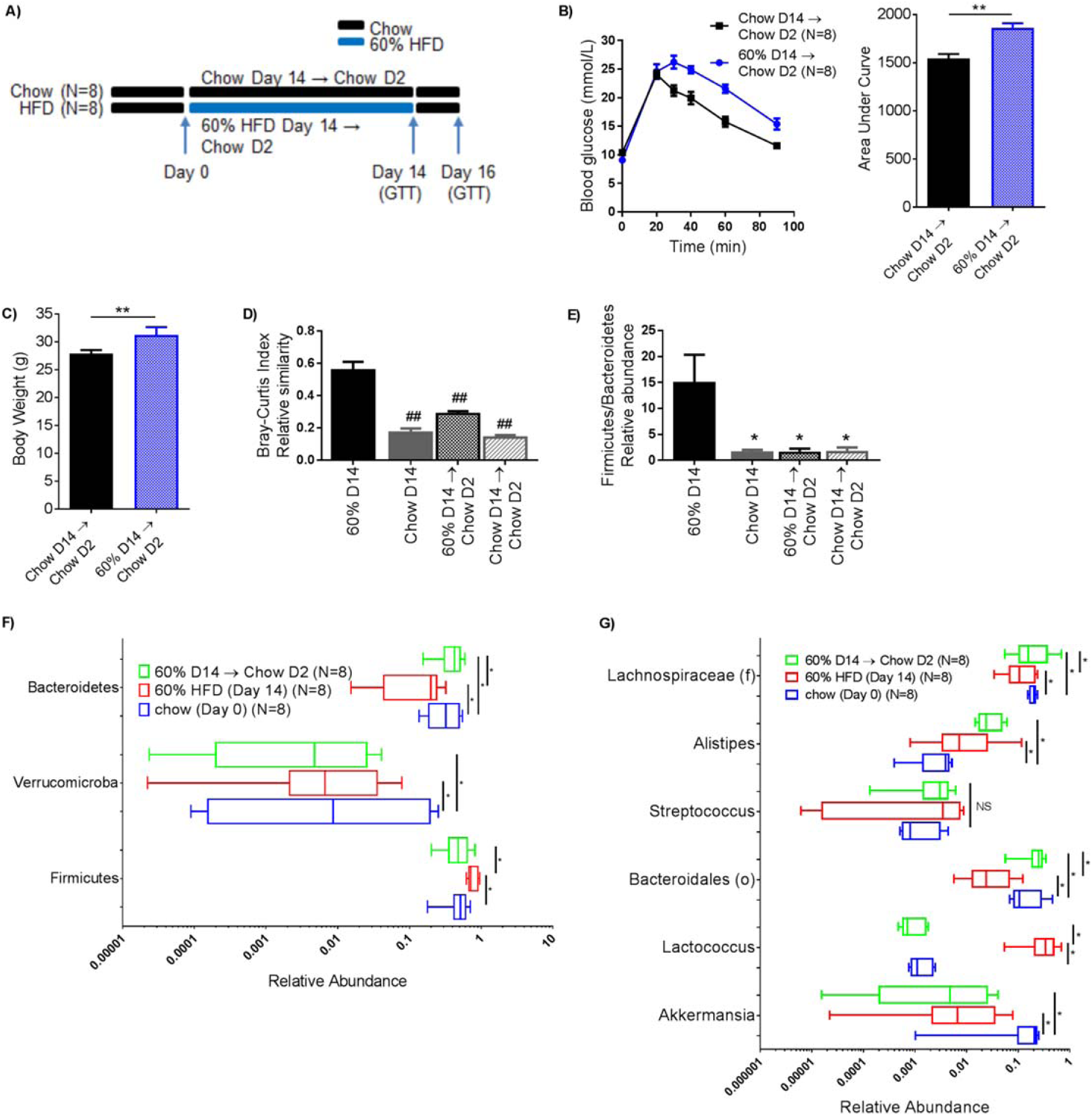
Glucose intolerance cannot be reversed by 2 days of chow diet following 14 days of HFD feeding. Mice were fed chow or 60% HFD (N= 8, 8) for 14 days before HFD removal for 2 days (Day 16) - schematic (A). B) GTT curve with AUC (2 g/Kg glucose, *i.p.*) and C) body mass on day 16. D) Bray-Curtis dissimilarity index relative to Day 14 on HFD. E) Firmicutes/Bacteroidetes abundance relative to Day 14 HFD. Phylum (F) and Genus (G) level changes in the microbiome on Day 14 (60% HFD) and Day 16 (2 day HFD removal). Statistical significance was measured as p<0.05 using Student t-test (B-C) or ANOVA (D, E) (* p<0.05; ** p<0.01; # p<0.001; ## p<.0001). For F and G ANOVA was used and false discovery rate was accounted for by controlling the familywise error rate using the Bonferroni correction; significance was accepted at p<0.05.

### Antibiotics attenuate glucose intolerance caused by long term, but not short term HFD

We next used antibiotics to mitigate diet-induced changes in the microbiota in order to test if short-term changes in microbes correspond with altered glycemia. We have previously established an antibiotic cocktail that causes profound changes in the gut microbiota and attenuates insulin resistance after prolonged HFD-feeding^17^, which has also been described by others^11^. The conditions for this experiment were based on our results showing that 4 days of HFD-feeding is sufficient to cause glucose intolerance (Fig. 1B). Mice were treated with antibiotics (0.5 g/L neomycin and 1.0 g/L ampicillin in the drinking water), which commenced 3 days prior to feeding a 60% HFD and continued for 4 days of HFD feeding (Fig 4A). This 7 day antibiotic treatment did not prevent glucose intolerance (Fig 4B) or hyperglycemia (Fig 4C) induced by feeding a 60% HFD for 4 days. Despite a small reduction in body mass (Fig 4C), antibiotics did not prevent increased adiposity or cause a change in fat mass during this short-term HFD-feeding (Fig. 4D). We next used this same antibiotic regimen to change the microbiota of mice fed a HFD for a longer period of time. Mice were fed a 60% HFD for 13 weeks followed by 7 days with or without antibiotics (Fig 4E). HFD-fed mice that received antibiotics showed improved glucose tolerance (Fig 4F) and lower fasting blood without changes in body mass (Fig. 4G). Overall, these results show that changing the microbiota with a specific antibiotic cocktail attenuated glucose intolerance and hyperglycemia after long-term, but not short-term HFD-feeding.

**Figure 4:**
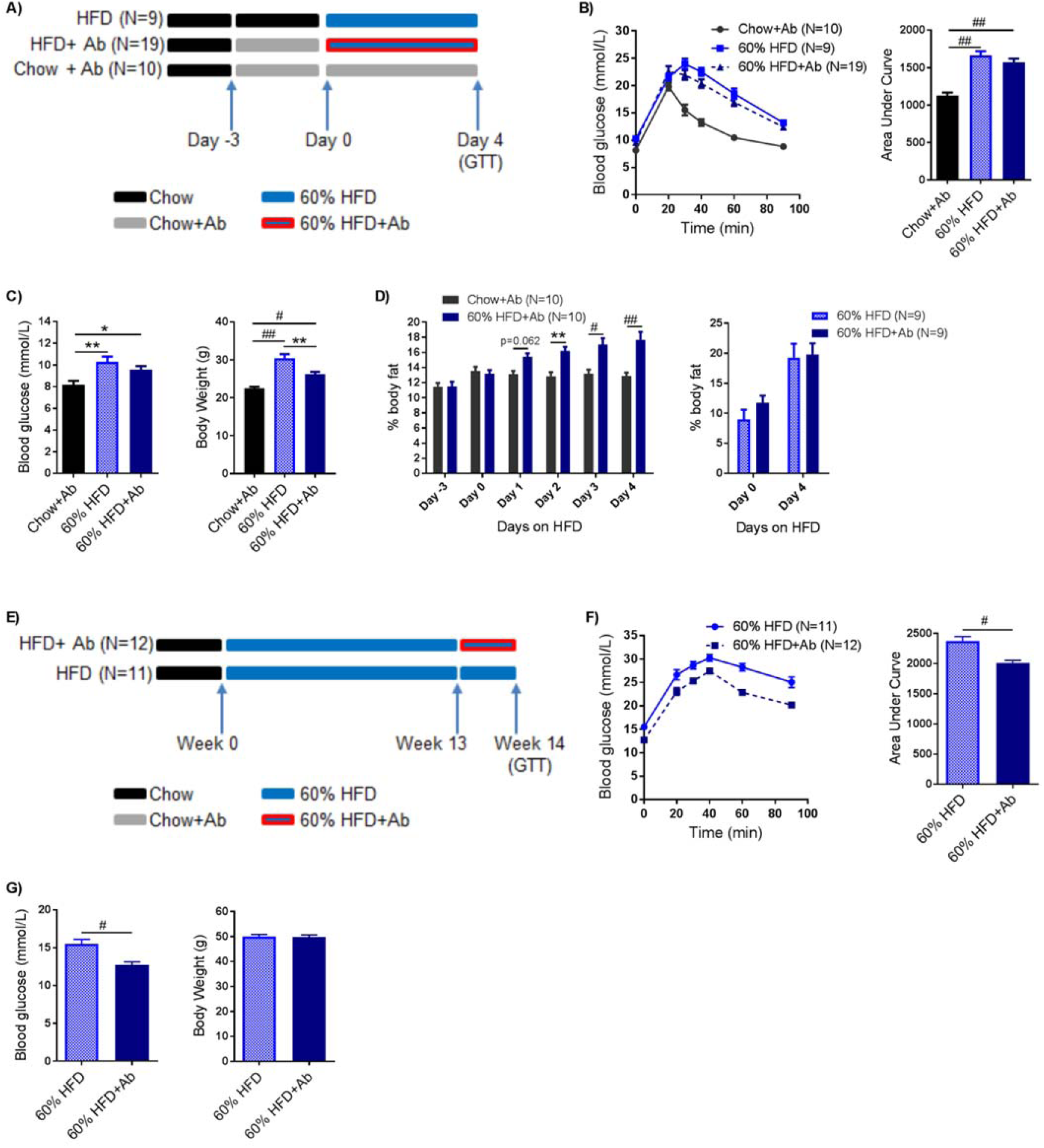
Antibiotics improve glucose tolerance of mice fed long-term, but not short-term HFD. A) Mice were treated with or without antibiotics (1mg/mL ampicillin and 0.5 mg/mL neomycin) in the water for 3 days before being placed on chow or 60% HFD with or without antibiotics for an additional 4 days (Chow+Ab=10; HFD=9; HFD+Ab=19). These mice were then tested for glucose tolerance (2 g/Kg glucose *i.p.*). GTT and AUC (B), fasting blood glucose and body mass (C) and body fat *%* (D) were measured. E) After 13 weeks of HFD, mice were treated with (N=12) or without (N=11) antibiotics (1mg/mL ampicillin and 0.5 mg/mL neomycin) in the drinking water for 1 week during the HFD and glucose tolerance (GTT) and AUC (F) and fasting blood glucose and body mass (G) were measured. Statistical significance was measured as p<0.05 using one-way ANOVA (B, C), Student t-test (F, G), or two-way ANOVA (D) with repeated measures (time). Post Hoc analysis was performed using Tukey’s multiple comparisons test (* p<0.05; ** p<0.01; # p<0.001; ## p<.0001).

### Microbiota from long-term, but not short-term HFD-feeding is sufficient to promote glucose intolerance

We next reconstituted germ-free mice to test if short-term HFD-induced changes in the microbiome are sufficient to promote dysglycemia. We first tested the cumulative effect of short-term diet-induced changes in microbiota over the first 6 weeks by continually exposing chow fed germ-free mice to the feces from donor mice fed a 60% HFD or chow diet (Fig 5A). Germ-free mice that received daily feces from 60% HFD mice or chow fed mice had similar glucose tolerance, fasting blood glucose, and percent body fat on day 4 (Fig 5C) and day 14 (data not shown) after microbiota reconstitution. However, after 45 days of microbiota reconstitution germ-free mice that received feces from 60% HFD mice were more glucose intolerant and had greater % body fat than those that received feces from chow fed mice (Fig. 5D). Phylum (Fig 5B) and genus (Fig 5E) level analysis shows microbial abundance in the feces of donor and recipient mice at day 45. Principle coordinate analysis (PCoA) and permutational analysis of variance (PERMANOVA - adonis) of Bray-Curtis dissimilarities show that changes in diet alter the beta diversity of the microbiota (Fig 5F). Specifically, ingesting a 60% HFD or receiving donor feces from mice fed this HFD altered the composition of the microbiota compared to a chow fed mouse (Fig 5F). Of note was the lack of transfer of *Lactococcus* from HFD donor mice to GF chow mice. This suggests that *Lactococcus* is present in the diet and not a major contributor to the observed metabolic phenotype. We also collected the feces from mice fed a 60% HFD for 4 days or a chow diet in order to specifically test repeated oral gavage of fecal slurry from this short term HFD. Germ-free mice that were gavaged (every 4 days) with the fecal slurry from HFD-fed or chow fed donor mice showed no difference in body mass, percentage body fat, glucose tolerance or insulin tolerance when tested on day 4 or day 25 after reconstitution (data not shown).

**Figure 5:**
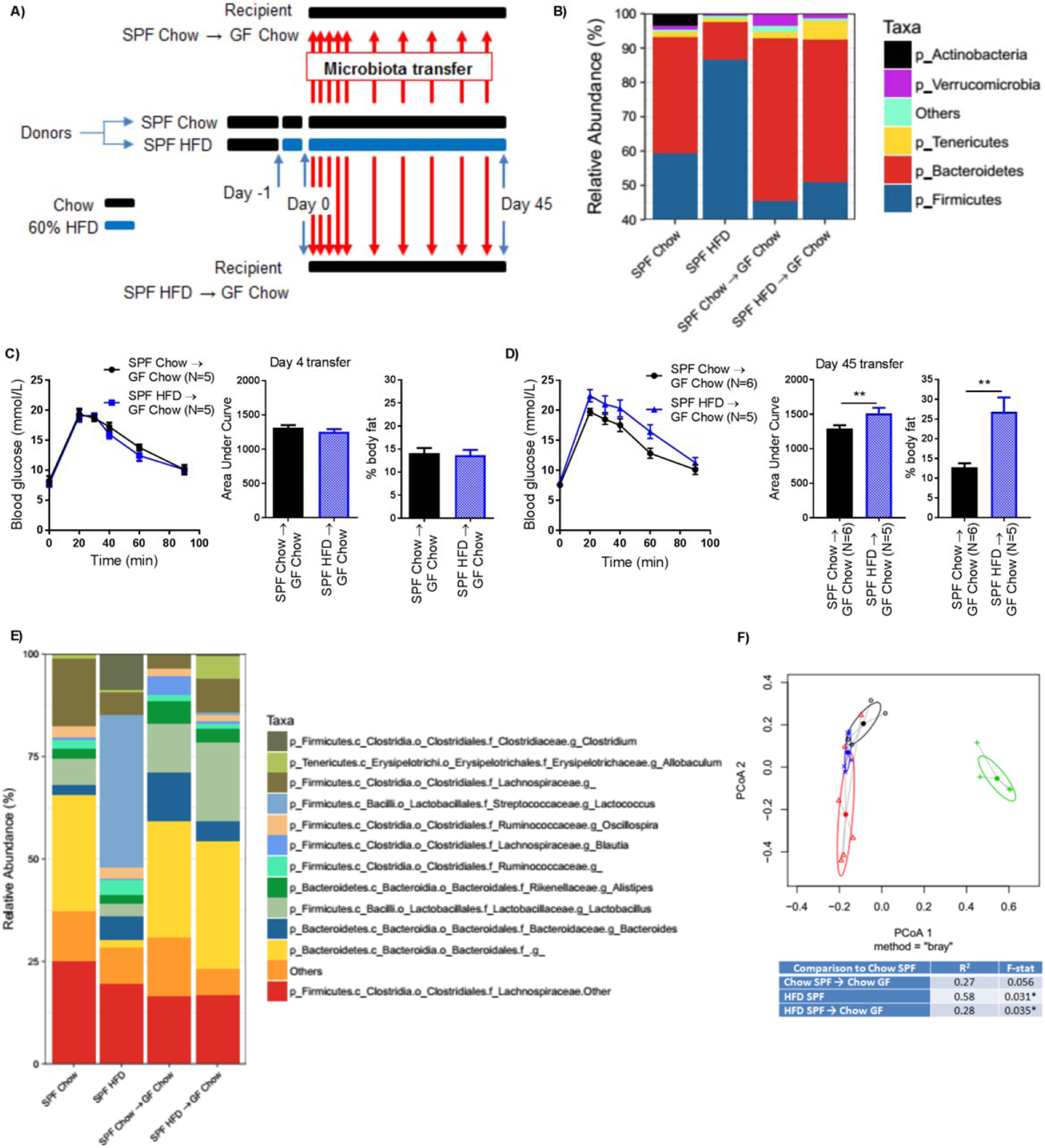
Gut microbiota from long-term, but not short-term HFD-fed mice causes glucose intolerance in germ-free mice. A) Schematic of experimental design. Donor mice were placed on chow or 60% HFD on Day -1. On Day 0, and each subsequent day, feces were transferred from donor chow or donor HFD cages to germ-free recipients fed chow diet. After 7 days feces were transferred once per week. B) Phylum level abundance of donor and germ free recipient feces after 45 days of microbiota transfer. On Day 4 (C) and Day 45 (D) of microbiota transfer (previously) germ-free, recipient mice were tested for glucose tolerance (N=5, 5). GTT, AUC, and *%* body fat are shown. Statistical significance was measured with a Student t-test (**p<0.01). E) Genus level abundance of donor and germ free recipient feces after 45 days of microbiota transfer. F) PCoA on Bray-Curtis dissimilarities (black = SPF Chow; green = SPF HFD; red = SPF Chow → GF Chow; blue = SPF HFD → GF Chow) and sources of variation due to each treatment assessed by PERMANOVA relative to the microbial profile in Chow donor mice (Day 45).

Finally, we reconstituted germ-free mice with the feces from donor mice that had been on a HFD for over 2 months and tested the effects on glycemia (Fig 6A). Mice that were used as microbiota donors were fed a chow or 60% HFD for 4 weeks prior to the experiment in order to discern if long term exposure of donor or recipient mice was the driver of changes in glycemia. Germ-free, recipient mice that were all fed a chow diet, but received daily feces from chow-fed or HFD-fed donors, had similar body mass, percentage body fat, and glucose tolerance when tested on day 4 (Fig 6C) or day 18 (data not shown) after reconstitution. However, recipient mice had increased glucose intolerance without changes in adiposity when tested 45 days after reconstitution with the microbiota from long term, HFD-fed donor mice (Fig. 6D). Phylum (Fig 6B) and genus (Fig 6E) level analysis shows microbial abundance in the feces of donor and recipient mice at day 45 when donor mice were on the HFD for 28 days prior to fecal transfer. PCoA and PERMANOVA (adonis) of Bray-Curtis dissimilarities again shows that ingesting a 60% HFD or receiving donor feces from mice fed a HFD altered the composition of the microbiota compared to a chow fed mouse (Fig 6F). These results show that the microbiota from long-term HFD feeding is sufficient to promote glucose intolerance when chow-fed mice are exposed to this dysbiosis for over a month (i.e. 45 days), but not for shorter duration such as 4 or 18 days.

**Figure 6:**
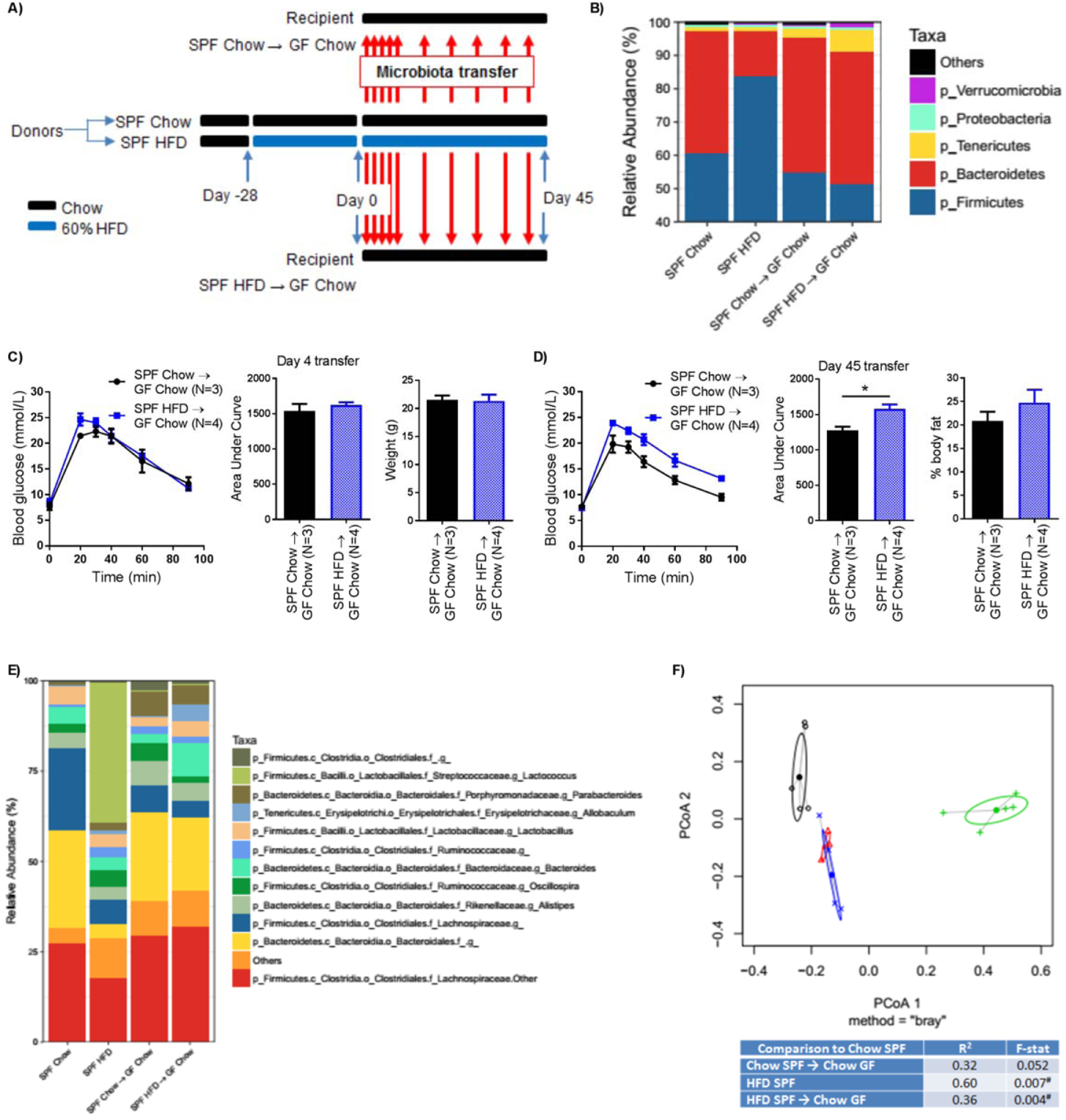
Long-term, but not short term HFD-induced dysbiosis is sufficient for transmissible glucose intolerance, independent of changes in adiposity. A) Schematic of experimental design. Donor mice were placed on chow or 60% HFD on Day -28. On Day 0, and each subsequent day, feces were transferred from donor chow or donor HFD cages to germ-free recipients fed chow diet. After 7 days feces were transferred once per week. B) Phylum level abundance of donor and germ free recipient feces after 45 days of microbiota transfer. On Day 4 (C) or Day 45 (D) of microbiota transfer germ-free recipient mice were tested for glucose tolerance (N=3, 4). GTT, AUC, and weight or % body fat are shown. Statistical significance was measured with a Student t-test (*p<0.05). E) Genus level abundance of donor and germ free recipient feces after 45 days of microbiota transfer. F) PCoA on Bray-Curtis dissimilarities (black = SPF Chow; green = SPF HFD; red = SPF Chow → GF Chow; blue = SPF HFD → GF Chow) and sources of variation due to each treatment assessed by PERMANOVA relative to the microbial profile in Chow donor mice (Day 45).

## Discussion

Environmental factors such as diet influence the risk of obesity, prediabetes and T2D. Obesity coincides with characteristics of prediabetes such as insulin resistance and glucose intolerance. The microbiota has emerged as a factor in obesity, but less is known about how the microbiota could connect the progression of obesity to the progression of prediabetes. Diet-induced dysbiosis is positioned to contribute to both obesity and glucose intolerance, but two ill-defined concepts were: 1) the timing of changes in the constituents of the microbiota relative to the onset of obesity and glucose intolerance and 2) whether dysbiosis contributes to dysglycemia via changes in adiposity during acute or chronic obesity.

We fed mice with two commonly used HFD and tracked the timing of taxonomic changes in the fecal microbiota versus the onset of glucose intolerance and changes in adiposity. It was known that HFD-feeding induces obesity and alters the gut microbiota. However, it was still not clear if diet-induced changes in the microbiota precede glucose intolerance or vice versa. We found that both a 45% and 60% HFD rapidly altered the constituents of the microbiota, which preceded overt changes in glucose tolerance in mice. Consistent with previous reports, we found that the fat content of the diet related directly to the magnitude of changes in microbial taxonomy^14^. We showed that these changes in the composition of the microbiota (and dietary fat content) directly related to the magnitude of glucose intolerance. We then showed that removal of the HFD, back to a chow diet (for only 2 days) caused a rapid reversion of the fecal microbiota such that many taxonomic markers resembled those of mice continually chow fed mice, an effect that is also consistent with previous findings in mice^14^. We extended these findings by showing that mice were still glucose intolerant after two days of replacing a HFD with a chow diet. We found that diet-induced changes in the constituents of the fecal microbiota precede and predict the magnitude of glucose intolerance in mice. Overall, our results are consistent with a model where the presence of a HFD is the major factor influencing changes in fecal microbiota composition, rather than insulin resistance altering the microbiota. We also showed that the first 3 days of a HFD resemble many of the changes seen after 14 weeks of a HFD regarding changes in the microbiota composition. These results initially supported the concept of using changes in the taxonomy of the microbiota as biomarkers of diet-induced glucose intolerance. However, the timing of divergent changes in the predicted microbial functional pathways such as metabolism and environmental information processing should be closely examined.

Subsequently, we used antibiotic treatment and reconstitution of germ-free mice to determine if diet-induced dysbiosis contributed to acute or chronic glucose intolerance. We also wanted to test if the rapid diet-induced dysbiosis caused features of prediabetes that were dependent on increased adiposity. It was already known that the constituents of the intestinal microbiota are altered in obesity and this dysbiosis can contribute to increased adiposity^8,9^. Our data showed that microbiota contributed to glucose intolerance, independent of changes in adiposity. In these experiments, we found that only long-term dysbiosis contributed to glucose intolerance. Despite rapid diet-induced changes in the constituents of the microbiota, we found no evidence of dysbiosis contributing to short-term HFD-induced glucose intolerance. Our results question the utility of using fecal microbial taxonomy as a biomarker for glucose intolerance during diet-induced obesity. Microbial function may be more important. We actually found that short term HFD feeding (for 3 days) altered the exact same predicted metagenomic annotations of metabolism and environmental information processing, which we have reported during long-term HFD feeding in mice^16^. However, short term HFD feeding affected these pathways in the opposite direction compared to long-term HFD feeding. Investigation of diet-induced changes in microbial function compared to taxonomy is warranted and it may be more useful to measure microbial-derived metabolites that can alter insulin resistance^18^.

Our results provide insight into the mechanism of dysbiosis-driven changes in glucose tolerance. It has previously been shown that metabolic inflammation contributes to insulin resistance and glucose intolerance during prolonged, but not short-term HFD-feeding^15^. Our results are consistent with microbiota contributing to glucose intolerance through metabolic inflammation during prolonged obesity. It is not clear how to reconcile our results in mice with recent results showing that 7 days of vancomycin or amoxcillin treatment had no impact on insulin sensitivity or substrate metabolism in obese humans^19^. The specific antibiotic used could be a key variable since (so far) we have found that only a combination of ampicillin and neomycin improves glucose tolerance, an effect not seen with either antibiotic alone or with vancomycin in HFD-fed mice (data not shown). Other groups have also shown that specific combinations of antibiotics improve glucose tolerance in obese mice^11,20^, but it is not yet clear if there is a difference between mice and humans.

The constituents of the microbiota are modifiable, which may provide therapeutic targets in obesity and prediabetes. Our results show that diet-induced dysbiosis can influence glucose tolerance independently of obesity. Therefore, microbiota-based interventions such as pre-and probiotics may be able to lower glucose intolerance and insulin resistance separately from obesity. Our results showed that the duration of diet-induced obesity is an important consideration in microbiota-targeted strategies aimed at attenuating dysglycemia versus adiposity.

## Methods

### Animal Experiments

All procedures were approved by McMaster University Animal Ethics Review Board. Specific pathogen free (SPF) C57BL/6 mice were born at McMaster University. Littermate mice were randomly placed on a HFD, where either 45% or 60% of calories are derived from fat (Research Diets, D12451 and D12492) or a chow diet (~5% fat). Blood glucose measurements and glucose and insulin tolerance tests were done after 6 h of fasting, as described ^17,21^. Body fat composition was measured using whole body echo-MRI (Bruker Minispec LF90-II). Antibiotics (1.0 mg/mL ampicillin and 0.5 mg/mL neomycin) were provided in the drinking water and changed every 2 days. Germ-free C57BL/6 mice were obtained from the Farncombe Gnotobiotic Unit of McMaster University and at 8-10 weeks of age. Germ-free mice were immediately and continually colonized by housing mice in soiled litter from SPF C57BL/6 donor mice. Where indicated, germ-free mice were given oral gavage every 4 days with a fecal slurry obtained from donor mice. Mice were housed using ventilated racks, and handled only in the level II biosafety hood to prevent bacterial contamination^17^.

### Bacterial profiling

Fecal samples were collected directly into sterile tubes and DNA was purified (Zymo Research Corporation: D6012), as described^17^. Illumina compatible PCR amplification of the variable 3 (V3) region of the 16S rRNA gene was completed on each sample. The Illumina MiSeq platform was used to sequence DNA products of this PCR amplification. A custom pipeline was used to process the FASTQ files, as previously described^17^. Operational Taxonomic Units (OTUs) were grouped using Abundant OTU+ based on 97% similarity. The 2011 version of the Greengenes reference database was used to assigned taxonomy to OTUs Ribosomal Database Project (RDP) classifier in Quantitative Insights Into Microbial Ecology (QIIME). QIIME and R scripts were used to calculate beta diversity using the Bray-Curtis dissimilarity and principal coordinate analysis, as previously described^22^. PERMANOVA were used to assess partitioning of variance in microbial communities with the adonis function from the vegan package in R^23^. Phylogenetic investigation of communities by reconstruction of unobserved states (PICRUSt) was used to predict the metagenome functional content from 16S rRNA gene data and group these into KEGG pathway maps, as described^16^. The LDA (Linear Discriminant Analysis) Effect Size (LEfSe) algorithm was used to identify differences in abundance of PICRUSt annotated pathways, where an LDA cut-off score of 2.0 and P value of 0.05 were used^25^. Sequencing characteristics are in Supplemental Table 1.

### Statistical analysis

An unpaired, two-tailed Student’s t-test was used to compare two groups. ANOVA and Tukey’s post hoc analysis was used to compare more than two means. Subsequently, false discovery rate (FDR) was accounted for via implementation of the Benjamini-Hochberg multiple testing adjustment procedure using R, where FDR-corrected p-values were estimated for all taxonomic and PICRUSt data. After adjustment for FDR, statistical significance was accepted at p < 0.05.

## Author Contributions

KPF researched the data, contributed to the discussion, and edited the manuscript. ED, BMD, RC researched the data. JCS analyzed data and contributed to discussion. JDS researched the data, derived the hypothesis, wrote the manuscript and is the guarantor of this work.

---

**Supplemental Figure 1:**
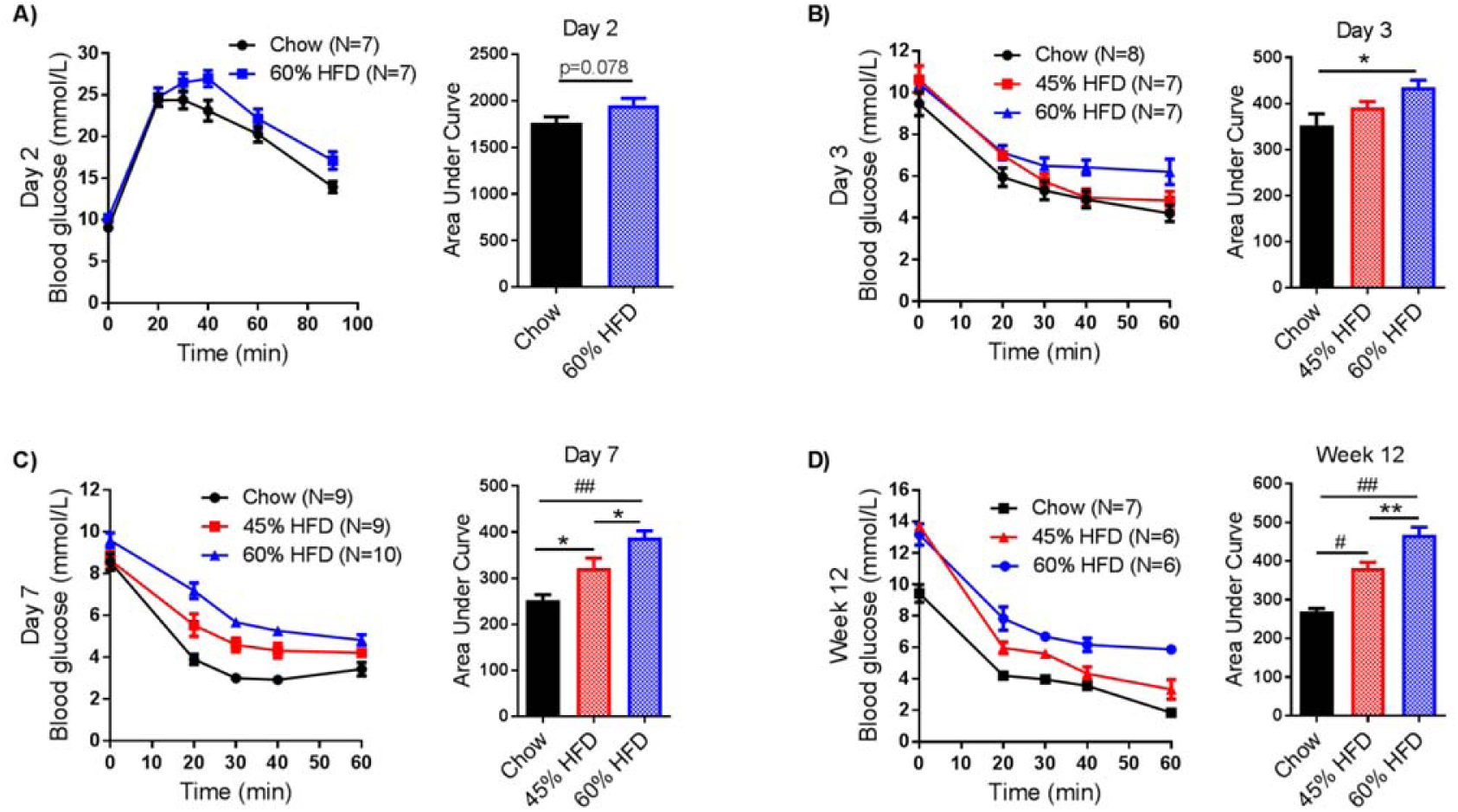
Short term high fat feeding causes insulin intolerance in mice. Mice were fed a chow, 45% HFD, or 60% HFD for 2 days (A: N=7, N/A, 7), 3 days (B: N=8, 7, 7), 7 days (C: N=9, 9, 10), or 12 weeks (D: N=7, 6, 6) before being tested for glucose (A) or insulin (B-D) tolerance *(i.p.)* with doses of 2g/Kg (A), 1.2 U/Kg (B), 0.9 U/Kg (C), or 1 U/Kg (D). Blood glucose measures were taken at indicated time points. Each figure shows the GTT/ITT curve and AUC. Statistical significance was measured as p<0.05 using one-way ANOVA. Post Hoc analysis was performed using Tukey’s multiple comparisons test (* p<0.05; ** p<0.01; # p<0.001; ## p<.0001).

**Supplemental Figure 2:**
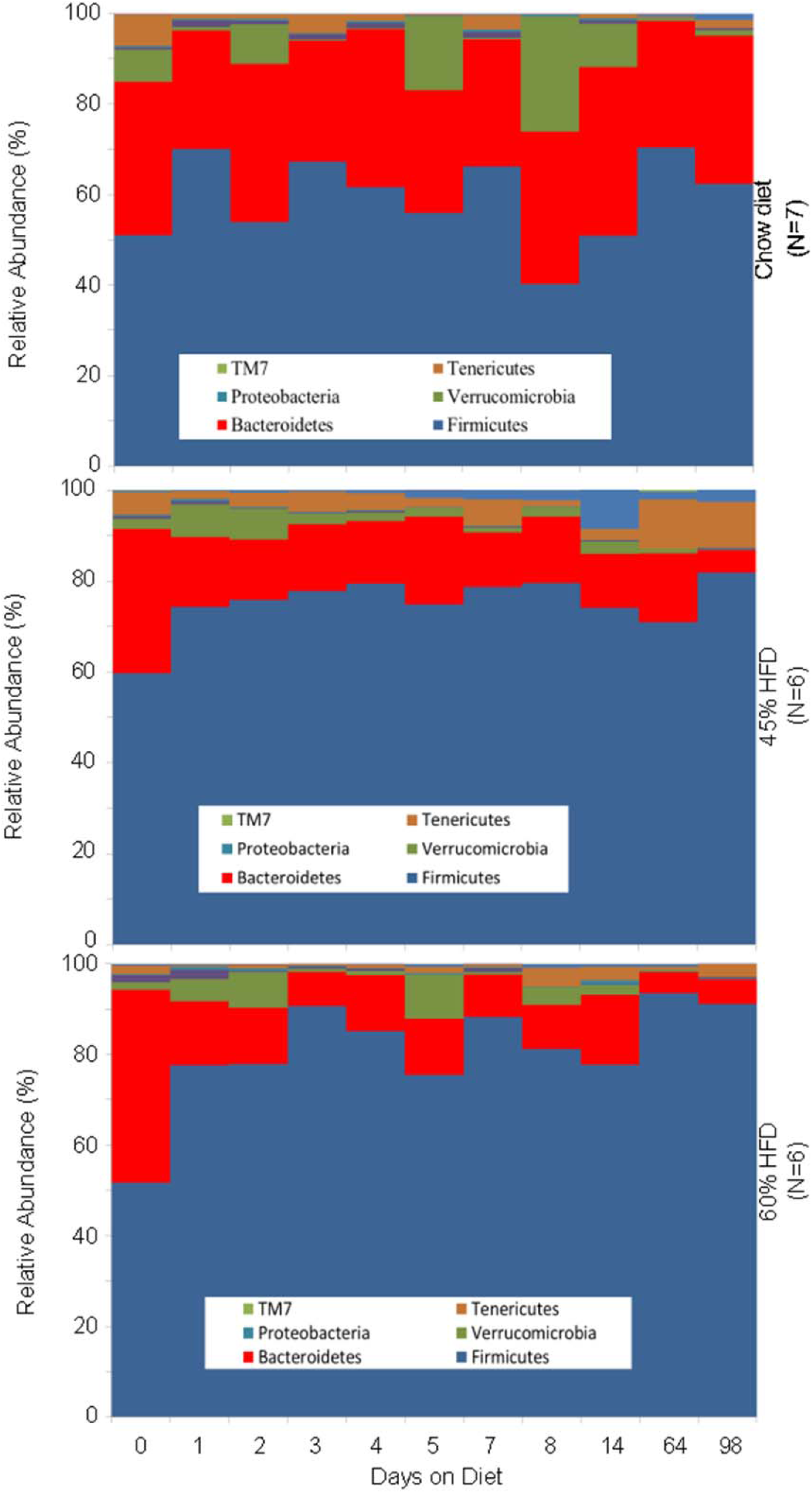
High fat feeding induces rapid changes in taxonomic composition of the fecal microbiome that remain stable over long-term feeding. Phylum level changes in mouse feces samples collected from onset of chow (N=7), 45% (N=6), and 60% (N=6) HFD up to 14 weeks after the start of high fat feeding.

**Supplemental Figure 3:**
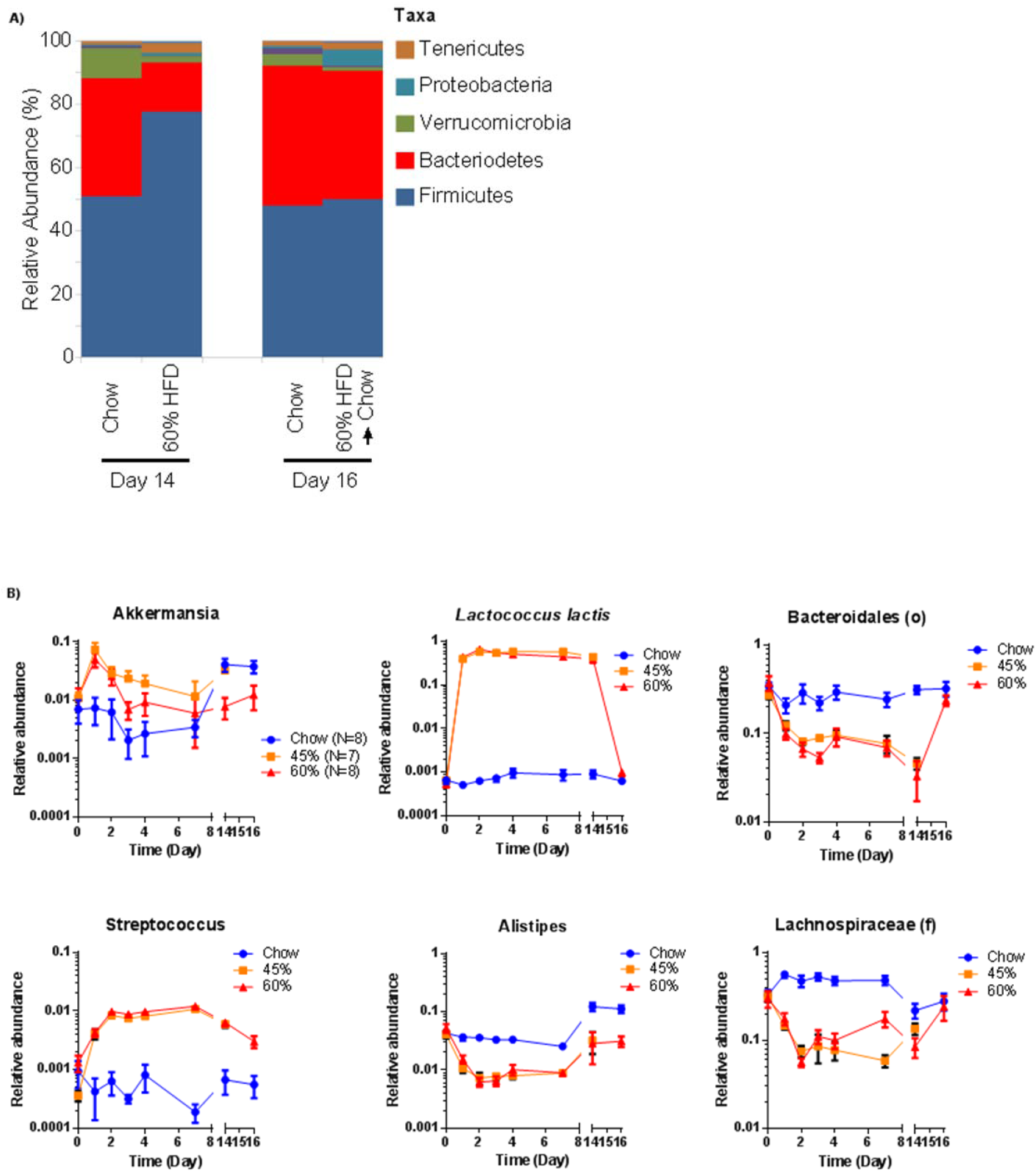
Dynamic changes in taxonomic composition of the fecal microbiome of the onset and removal of a HFD in mice. A) Phylum level abundance on Day 14 versus Day 16 of mice fed a chow diet or 60% HFD before (Day 14) or after (Day 16) 2 days of changing this 60% HFD back to a chow diet (N = 8, 8). B) Relative abundance changes over 14 days of high fat feeding/2 days of diet removal back to a chow diet (N = 8, 7, 8) for *Akkermansia, Lactococcus lactis*, Bacteriodales (o), *Streptococcus, Alistipes* and Lachnospiraceae (f). Data are mean + SEM.

## References

1. Nathan, D. M. et al. Impaired fasting glucose and impaired glucose tolerance: implications for care. Diabetes Care 30, 753–759 (2007).

2. Schellenberg, E. S., Dryden, D. M., Vandermeer, B., Ha, C. & Korownyk, C. Lifestyle interventions for patients with and at risk for type 2 diabetes: a systematic review and meta-analysis. Ann. Intern. Med. 159, 543–551 (2013).

3. Zeevi, D. et al. Personalized Nutrition by Prediction of Glycemic Responses. Cell 163, 1079–1094 (2015).

4. Ley, R. E. et al. Obesity alters gut microbial ecology. Proc. Natl. Acad. Sci. U. S. A. 102, 11070–11075 (2005).

5. Qin, J. et al. A metagenome-wide association study of gut microbiota in type 2 diabetes. Nature 490, 55–60 (2012).

6. Forslund, K. et al. Disentangling type 2 diabetes and metformin treatment signatures in the human gut microbiota. Nature 528, 262–266 (2015).

7. Bäckhed, F. et al. The gut microbiota as an environmental factor that regulates fat storage. Proc. Natl. Acad. Sci. U. S. A. 101, 15718–15723 (2004).

8. Ridaura, V. K. et al. Gut microbiota from twins discordant for obesity modulate metabolism in mice. Science 341, 1241214 (2013).

9. Turnbaugh, P. J. et al. An obesity-associated gut microbiome with increased capacity for energy harvest. Nature 444, 1027–1031 (2006).

10. Pedersen, H. K. et al. Human gut microbes impact host serum metabolome and insulin sensitivity. Nature (2016). doi:10.1038/nature18646

11. Cani, P. D. et al. Changes in gut microbiota control metabolic endotoxemia-induced inflammation in high-fat diet-induced obesity and diabetes in mice. Diabetes 57, 1470–1481 (2008).

12. Schertzer, J. D. et al. NOD1 activators link innate immunity to insulin resistance. Diabetes 60, 2206–2215 (2011).

13. Vrieze, A. et al. Transfer of intestinal microbiota from lean donors increases insulin sensitivity in individuals with metabolic syndrome. Gastroenterology 143, 913–916.e7 (2012).

14. Carmody, R. N. et al. Diet dominates host genotype in shaping the murine gut microbiota. Cell Host Microbe 17, 72–84 (2015).

15. Lee, Y. S. et al. Inflammation is necessary for long-term but not short-term high-fat diet-induced insulin resistance. Diabetes 60, 2474–2483 (2011).

16. Denou, E., Marcinko, K., Surette, M. G., Steinberg, G. R. & Schertzer, J. D. High-intensity exercise training increases the diversity and metabolic capacity of the mouse distal gut microbiota during diet-induced obesity. Am. J. Physiol. Endocrinol. Metab. 310, E982–993 (2016).

17. Denou, E. et al. Defective NOD2 peptidoglycan sensing promotes diet-induced inflammation, dysbiosis, and insulin resistance. EMBO Mol. Med. 7, 259–274 (2015).

18. Perry, R. J. et al. Acetate mediates a microbiome-brain-β-cell axis to promote metabolic syndrome. Nature 534, 213–217 (2016).

19. Reijnders, D. et al. Effects of Gut Microbiota Manipulation by Antibiotics on Host Metabolism in Obese Humans: A Randomized Double-Blind Placebo-Controlled Trial. Cell Metab. 24, 63–74 (2016).

20. Membrez, M. et al. Gut microbiota modulation with norfloxacin and ampicillin enhances glucose tolerance in mice. FASEB J. Off. Publ. Fed. Am. Soc. Exp. Biol. 22, 2416–2426 (2008).

21. Schertzer, J. D. et al. A transgenic mouse model to study glucose transporter 4myc regulation in skeletal muscle. Endocrinology 150, 1935–1940 (2009).

22. Caporaso, J. G. et al. QIIME allows analysis of high-throughput community sequencing data. Nat. Methods 7, 335–336 (2010).

23. Oksanen, J et al. vegan: Community Ecology Package. (2016).

24. Langille, M. G. I. et al. Predictive functional profiling of microbial communities using 16S rRNA marker gene sequences. Nat. Biotechnol. 31, 814–821 (2013).

25. Segata, N. et al. Metagenomic biomarker discovery and explanation. Genome Biol. 12, R60 (2011).

